# MycoCirc: A Pan-Fungal Multi-Modal Pretrained Model for Fungal circRNA Prediction from Genome Sequence

**DOI:** 10.64898/2026.06.29.735431

**Authors:** Xueyan Hu, Yueqi Jin, Juan Wang, Ence Yang

**Affiliations:** Department of Microbiology and Infectious Disease Center, Beijing Key Laboratory for Research and Translation of Immunological and Molecular Diagnostic Technologies for Major Viral Infectious Diseases, School of Basic Medical Sciences, Peking University Health Science Center, Beijing 100191, China; Department of Radiation Medicine, School of Basic Medical Sciences, Peking University Health Science Center, Beijing 100191, China; Institute of Systems Biomedicine, School of Basic Medical Sciences, Peking University Health Science Center, Beijing 100191, China

**Keywords:** Fungal circRNA, CircRNA prediction, Fungal genomics, Multi-modal learning, Transfer learning

## Abstract

**Motivation:** Exploring the fungal circular RNA (circRNA) landscape is bottlenecked by both experimental and computational limits. While standard mRNA-seq systematically discards circRNAs due to their lack of poly(A) tails, high-cost total RNA-seq remains prohibitive for large-scale screening. Consequently, discovery relies heavily on computational prediction. However, existing models trained exclusively on human or plant sequences fail in fungi because of the extreme genomic divergence across fungal lineages, which span from intron-poor *Candida* to intron-rich filamentous fungi. As a result, no computational framework currently exists for de novo fungal circRNA prediction, leaving the vast majority of non-model fungi entirely inaccessible.

**Results:** We present mycoCirc, the first end-to-end pan-fungal multi-modal pretrained model for fungal circRNA prediction, integrating three mandatory modalities with bidirectional cross-attention for donor–acceptor site interaction. Pre-trained on 22 strains with 16,483 positive gene–circRNA associations spanning Ascomycete yeast, Basidiomycete yeast, and Filamentous fungi groups and fine-tuned per group using 5-fold cross-validation, mycoCirc achieved AUROC 0.69–0.70 on held-out test species under Mode A (Genome+GTF, no RNA-seq), substantially outperforming JEDI (0.51–0.57) and CircPCBL (0.49–0.53). Cross-species evaluation on four independent fungi datasets demonstrated robust generalization across all fine-tuned variants (AUROC 0.63–0.72). Beyond gene-level classification, the JunctionEncoder module enabled backsplicing junction identification for detailed circRNA validation. We further build mycoCircAtlas, a companion database providing 319,860 high-confidence gene–circRNA predictions across 768 fungal species from Ensembl Fungi Release 113, enabling researchers to query precomputed predictions and design validation primers without local model deployment.

**Availability and Implementation:** mycoCirc is freely available under the MIT license at https://github.com/yukkikou/mycoCirc and https://doi.org/10.5281/zenodo.20823962. The mycoCircAtlas database is available at http://116.62.58.192.

## Introduction

Circular RNAs (circRNAs) are a class of covalently closed long non-coding RNAs generated through backsplicing, a non-canonical splicing event in which a downstream 5′ splice site joins to an upstream 3′ splice site^1^. Initially considered splicing artefacts, circRNAs are now recognised as functionally important molecules involved in microRNA sponging, transcriptional regulation, and protein scaffolding^2,3^. In fungi, circRNAs have been implicated in stress responses, pathogenicity, developmental transitions, and regulated morphological switching^4–9^. However, circRNA research is limited to a small number of fungi, and information on circRNAs remains scarce for the vast majority of fungi species.

The pervasive gap in fungal circRNA research appears to persist primarily due to severe technical constraints in data acquisition. Fungal RNA-seq experiments often present formidable challenges because many pathogenic species require Biosafety Level 2 or 3 facilities^10^, and RNA extraction from filamentous mycelia frequently yields constrained quantities due to their rigid, chitin-rich cell walls and complex secondary metabolites^11,12^. Crucially, because circular RNAs lack polyadenylated tails, conventional mRNA-seq approaches utilizing poly(A) enrichment are entirely unsuitable for circRNA discovery as they systematically discard circular transcripts during library preparation^13^. Consequently, researchers must rely on total RNA-seq, a methodology wherein ribosomal RNA characteristically constitutes the vast majority of the total yield. To uncover low-abundance circRNA signals, efficient ribosome depletion is mandatory, yet conventional depletion strategies optimised for mammalian systems frequently show limited efficacy on fungal RNA. This technical bottleneck necessitates specialised, fungi-specific procedures that remain relatively scarce for most non-model fungi^14^. Characterising the circRNA landscape across the fungal kingdom remains therefore experimentally prohibitive, establishing a critical need for computational frameworks capable of high-throughput prediction directly from genomic sequences without relying on transcriptomic evidence.

However, executing genome-only prediction in fungi introduces distinct computational complexities driven by their unique genomic architectures. In mammalian systems, the vast majority of circRNAs arise from multi-exon genes through canonical exon skipping, a process heavily facilitated by complementary repeats within flanking introns^15^. In contrast, the proportion of intron-rich genes varies widely across fungal clades, where, for instance, a substantial majority of genes in certain *Candida* species possess single-exon architectures^16,17^. Given that single-exon circRNAs form through self-circularization rather than standard exon-intron splicing, the predictive rules derived from multi-exon backsplicing are fundamentally inapplicable to these unique fungal contexts^18,19^. While current benchmarks, such as JEDI and CircPCBL, achieve sequence-only prediction in their respective domains, they rely exclusively on raw nucleotide sequences to capture multi-exon splicing signals^20,21^. As they completely omit structural gene annotation features, these models lack the capacity to accommodate the heterogeneous mixture of single-exon and multi-exon profiles characteristic of fungal genomes, leaving non-model fungi difficult to scan effectively.

Here, we introduce mycoCirc, an end-to-end multi-modal pretrained model for pan-fungal circRNA prediction, trained on circRNAs identified from real rRNA-depleted RNA-seq datasets across 22 fungal strains spanning three taxonomic groups. Unlike existing single-modal sequence-based methods, mycoCirc integrates three mandatory input modalities (gene annotation, genomic sequence context, and junction-level sequence features) through a hierarchical multi-modal fusion architecture, establishing a pretraining with per-group fine-tuning paradigm that enables robust cross-species generalisation. Critically, mycoCirc operates entirely on reference genomes and gene annotations without requiring external RNA-seq data at inference, thereby providing an accessible solution for circRNA screening in data-scarce non-model fungal species.

## System and Methods

### In-house Sequencing and Species Taxonomy

To build a robust foundation for pan-fungal analysis, we constructed a comprehensive pan-fungal circRNA dataset spanning 22 fungal strains of 19 fungi species, harnessing our newly sequenced, in-house rRNA-depleted strand-specific RNA-seq data generated in biological triplicates as previously described using species-specific rRNA-depletion probes^14^. To define these groupings robustly, single-copy protein-coding genes were identified via BUSCO and subsequently concatenated into a supermatrix for phylogenetic tree reconstruction using RAxML (Supplemental Figure 1) ^22,23^. The dataset were structurally categorized into three major taxonomic groups, comprising 6 Ascomycete yeast species (*Candida albicans*, 2 strains of *C. tropicalis, C. glabrata, C. auris, Pichia kudriavzevii*, and *Saccharomyces cerevisiae*), 5 Basidiomycete yeast species (2 strains of *C. neoformans var. grubii, C. neoformans* var. *neoformans, C. floricola, 2 strains of C. gattii*, and *C. laurentii*), and 8 filamentous fungi species (*Aspergillus fumigatus, A. nidulans, Fusarium dimerum, F. proliferatum, F. venenatum, Neurospora crassa, Penicillium chrysogenum*, and *Schizosaccharomyces pombe*). Notably, The Ascomycete yeast group includes S. cerevisiae due to its phylogenetic proximity and comparable genome architecture, representing a functional grouping rather than a strict taxonomic classification^24^. An analogous rationale applies to the Filamentous fungi group, where *S. pombe* is strategically integrated to capture conserved eukaryotic processing mechanisms despite traditional taxonomic divergence^25^.

All baseline gene expression data utilized for model training were newly generated in-house via high-quality, strand-specific, paired-end 150 bp (PE150) RNA-seq sequenced on the Illumina platform in biological triplicates, using HISAT2 and StringTie2^26^. To ensure the maximum reliability of circRNAs utilized for model training, candidate circRNAs were initially identified from our in-house RNA-seq data using three independent detection pipelines, including DCC^27^, CIRI3^28^, and CLEAR3^29^. Previous comparative studies have consistently demonstrated that these alignment-based detection tools inherently possess high specificity^30^. Then, the resulting circRNA predictions were integrated and quantified via CIRIquant to yield high-confidence, high-quality circRNA profiles^31^. Finally, each strain is associated with a reference genome FASTA, a gene annotation GTF from Ensembl Fungi (release113), gene expression data, and circRNA data from CIRIquant. After filtering to retain only exon and intron circRNA types and excluding *A. nidulans* (which had few circRNA records), the training set comprised 16,483 positive gene–circRNA associations across 17 training strains and 4 test strains (Supplemental Table 1).

### Held-Out Test Strategy and Negative Sample Construction

Each taxonomic groups designated one species as a held-out test species that is never seen during training or fine-tuning. These are *C. auris* for the Ascomycete yeast group, *C. neoformans* var. *neoformans* for the Basidiomycete yeast group, and *F. venenatum* for the Filamentous fungi group. This leave-one-species-out evaluation provides a stringent test of cross-species generalisation. Additionally, the model’s true zero-shot performance was validated using external data from four completely unseen species, including *Aspergillus cristatus, Trichophyton mentagrophytes, Trichophyton rubrum*, and *Talaromyces marneffei* (Supplemental Table 1) ^4,7–9^. These independent cohorts, derived from previously published studies on fungal circRNA identification, provide a rigorous foundation for cross-species evaluation as they were entirely omitted from the training pipeline. For species with available metadata, circRNAs were retrieved directly from their published reference lists, whereas those lacking metadata were characterized using the aforementioned circRNA identification pipeline. For each strain, positive genes are defined as those with at least one exon or intron circRNA record. Negative samples are selected from the same strain’s genes that have no circRNA record, with gene length distribution matched to the positive set, and a negative-to-positive ratio of 1:1.

### Multi-Modal Input Representation

The mycoCirc framework incorporates five input modalities to accommodate different stages of the machine learning pipeline. The three core modalities required during pre-training consist of gene annotation features from gene annotation features (GTF), genomic sequence contexts (Genome), and junction-level flanking sequences derived from identified circRNAs. Conversely, the two optional modalities, namely circRNA expression and gene expression, are utilized exclusively during the fine-tuning phase. This distinction between core and optional modalities is pivotal for model deployment, as only the three core modalities are required at the inference stage, eliminating the need for transcriptomic data.

#### Gene annotation (GTF) modality

For each gene, we extracted a 17-dimensional feature vector from the GTF annotation. The first 8 dimensions were numeric gene-structure features: exon count, log-transformed total exon length, log-transformed total intron length, log-transformed CDS length, exon density (exons per kb), relative gene length (normalised to 10 kb genes), and binary indicators for multi-exon status and CDS presence. The 9th dimension was a gene biotype identifier subsequently embedded to 8 dimensions inside the GTFEncoder, and the remaining 8 dimensions were reserved for future feature expansion. These features encoded the architectural information most relevant to circRNA biogenesis, capturing the intuition that genes with more exons and longer introns have more backsplicing opportunities.

#### Genomic context modality

A symmetric ±5 kb window centred on each gene was profiled using sliding 50 bp bins to compute eight sequence-composition statistics per bin, including GC content, the four mononucleotide frequencies (A, C, G, T), the observed-to-expected CpG ratio, the AT/GC ratio, and the Shannon entropy, producing a 200 × 8context tensor that captures the local genomic environment.

#### Junction modality

For each exon boundary, we extracted 300 bp flanking sequences (150 bp upstream plus 150 bp downstream) centred on the splice site, which was identified by hyperparameter sweeping. These flanks were encoded through three parallel pathways whose outputs were fused and processed by bidirectional cross-attention between donor and acceptor sites.

#### Species embedding

A 32-dimensional species representation was constructed by combining a learned species lookup table with an 8-dimensional phylogenetic embedding derived from classical multidimensional scaling of a BUSCO-based RAxML phylogenetic distance matrix across the training strains. The two components were concatenated and passed through a linear–LayerNorm–ReLU block to produce the final 32-dimensional species vector, providing taxonomic context that enabled the model to adjust predictions based on species-specific genome architectures.

#### Expression modality (fine-tuning only)

log1p-transformed and z-score normalised CircExp (circRNA expression) and GeneExp (gene expression) values were encoded through two parallel MLPs (one per channel), fused via a two-head cross-attention module with a feed-forward refinement, and projected to the fusion dimension. This modality was not required at inference time.

### Model Architecture

MycoCirc employed a hierarchical multi-modal architecture totalling 775,858 parameters, organised into seven components.

#### GenomicContextEncoder

The GenomicContextEncoder processed the 200 × 8genomic profile through a three-layer dilated convolutional network dilations[1,2,4], 64 filters, kernel size 7), a single-layer bidirectional GRU (hidden size 64), and learned attention pooling, producing a 128-dimensional context vector **v**_genome_.

#### GTFEncoder

The GTFEncoder split the 17-dimensional GTF feature vector into a 16-dimensional numeric block (8 active features plus 8 reserved slots, with the biotype identifier excluded) and a scalar biotype index. The biotype index was mapped to an 8-dimensional embedding, concatenated with the numeric block, and projected by a two-layer MLP (24 → 64 → 128( with LayerNorm and ReLU after each layer, producing **v**_gtf_ ∈ ℝ^128^.

#### JunctionEncoder

The JunctionEncoder processed all exon boundaries of a gene collectively through three parallel encoding pathways whose outputs were fused and then refined through bidirectional cross-attention.

### *K*-mer embedding pathway

Flanking sequences were tokenised into 3-mers, embedded, and processed by a bidirectional GRU (*h*_gru_=64, bidirectional) with learned attention pooling to produce **s**_JEDI_ ∈ ℝ^128^per site.

### One-hot CNN-BiGRU pathway

The raw flanking sequence was one-hot encoded, processed by multi-scale convolutions (kernel sizes 3, 5, 7) and a bidirectional GRU to produce **s**_CNN_∈ ℝ^64^.

### *K*-mer frequency GLT pathway

k-mer frequencies *k*=1..4, 340-dim) were processed by a Group Linear Transform (*G*=4) to produce **S**_GLT_∈ ℝ^64^

The three pathway outputs were fused:

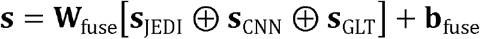

producing a 128-dimensional vector per site.

Bidirectional cross-attention. Let 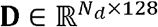and 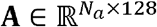 be donor and acceptor site matrices, respectively. Cross-attention produced context vectors followed by residual connections and attention pooling:

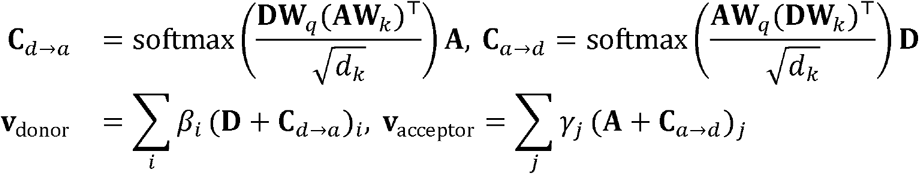

where **W**_*q*_,**W**_*k*_∈ ℝ^128 × 16^ *d*_*k*_=16, and *β γ* were learned attention weights. The pooled vectors were concatenated and projected to produce the final junction vector **j** ∈ ℝ^64^ . The cross-attention mechanism scored donor–acceptor compatibility, providing the signal that enabled backsplicing exon-pair identification.

#### Fusion Module

Each modality was linearly projected to a common dimension(**p**_(·)_∈ ℝ^128^)and fused through a two-layer MLP:

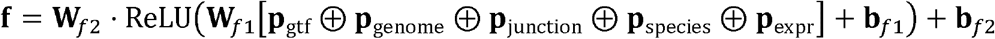

Where **W**_*f*1_∈ ℝ^640 × 256^, **W**_*f*2_∈ ℝ^256 × 128^,and **p**_expr_ was zeroed when expression data was absent.

#### Prediction Heads

MycoCirc produced two outputs from the fused representation.

Gene-level prediction. The GeneHead was a two-layer MLP (128 → 64 → 1)( with LayerNorm and ReLU on the hidden layer and a sigmoid on the output, producing the circRNA probability for each gene:

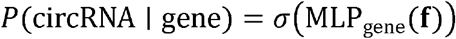

#### Junction-level prediction

The bidirectional cross-attention inside the JunctionEncoder yielded a raw logit matrix 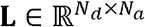 over all donor–acceptor 253 exon-pair candidates for a gene, where *N*_*d*_ and *N*_*a*_ were the numbers of donor and acceptor sites. These logits were flattened into a score vector of length *N*_*d*_ ·*N*_*a*_ and candidate pairs were ranked by descending score. At inference, the top-*K* scoring exon pairs were reported as the predicted backsplice junctions for a gene classified as circRNA-positive. The cross-attention logits thus served a dual role: they were supervised by the junction-level BCE loss during Stage 2 pre-training, and they directly powered the junction-prediction capability evaluated in the Results.

#### Inference modes

Two forward-pass modes were defined, differing in whether the expression modality was used. In *Mode A* (Genome+GTF, no expression data), the ExpressionEncoder was inactive and **p**_expr_ was zeroed, so the fused representation depended only on the four mandatory modalities (GTF, genomic context, junction, species); this mode required only a reference genome and gene annotation and constituted the primary deployment use case for understudied fungi. In *Mode B* (Genome+GTF+GeneExp), the ExpressionEncoder was activated and the host gene-expression channel (GeneExp) was fed through the fusion, while the circRNA-expression channel (CircExp) was zeroed to avoid leaking the circRNA signal at inference; this mode was applicable only when within-species RNA-seq data were available. Both modes produced gene-level probabilities via the GeneHead and junction-level rankings via the cross-attention logits.

### Pre-Training Protocol

Pre-training proceeded in two stages on all 17 training strains simultaneously. In Stage 1 (50 epochs), the GTFEncoder, GenomicContextEncoder, FusionModule, and GeneHead were trained with binary cross-entropy loss *ℒ*_1_, while the JunctionEncoder remained frozen; freezing the JunctionEncoder in Stage 1 prevented the unstable cross-attention layer from destabilising modality-specific feature learning. In Stage 2 (100 epochs), all modules were unfrozen and trained with a combined loss:

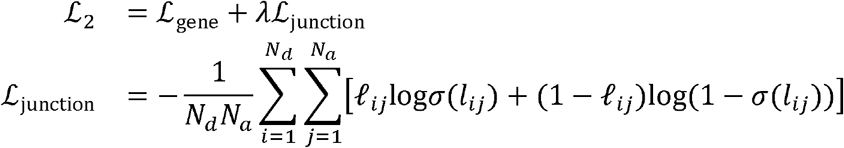

Where *l*_*ij*_ ∈{0,1} indicated whether exon pair (*i,j*) was a known backsplice junction and . This loss trained the cross-attention to assign higher scores to genuine backsplice pairs, though the extreme class imbalance (typically 1–2 positive pairs among hundreds of candidates) limited its effectiveness for ranking.

AdamW was used with learning rate10^−3^(Stage 1) and 5 ×10^−4^ (Stage 2), batch sizes 32 (Stage 1) and 16 (Stage 2), and weight decay10^−5^

### Fine-Tuning Protocol

Each taxonomic group was fine-tuned independently from the shared pre-trained backbone. For each group, we performed 5-fold cross-validation by strain. In each fold, *N* −1 training strains were fine-tuned and 1 held-out strain served as the validation set. The best fold (highest validation AUROC) was selected and evaluated on the group’s designated held-out test species. During fine-tuning, the ExpressionEncoder was activated and received CircExp and GeneExp as additional input features. We tested two inference modes. Mode A (Genome+GTF, no expression data) used the pre-train-style forward pass, and Mode B (Genome+GTF +GeneExp) used the fine-tune-style forward pass. Fine-tuning used AdamW (lr=5 ×10^−5^, batch size 16) with early stopping (patience 10 epochs) and a linear-probe strategy in which all encoder parameters were initially frozen except the prediction head and expression encoder, then optionally unfrozen for joint optimisation (maximum 100 epochs).

### Baseline Methods

We compared mycoCirc against JEDI and CircPCBL. JEDI was retrained from scratch for each panel using the training strains of the corresponding group. In contrast, CircPCBL used the same pretrained Plant model for all three panels, independent of group assignment. Both baselines were evaluated on the same held-out test species and external unseen cohorts defined above, using the same positive/negative gene sets and 1:1 length-matched sampling, to ensure direct comparability with mycoCirc.

### Implementation

MycoCirc was implemented in Python 3.9+ using PyTorch ≥ 2.0. The model was deployable on a single GPU with 8 GB memory and supported two use cases: training mode for users with circRNA metadata, and inference mode for users with only a reference genome and GTF. Pre-trained checkpoints for all three groups were provided. The companion mycoCircAtlas database was built by applying the group-fine-tuned mycoCirc models in Mode A to all 768 fungal species in Ensembl Fungi Release 113, and was deployed as a public web application (http://116.62.58.192), and an integrated Primer3 module for backsplice-junction primer design.

## Results

### Overview of mycoCirc

We adopt a five-modality fusion framework to construct a cross-species fungal circRNA prediction model, with GTF annotation, genomic sequence context, and junction flanking sequences as the core mandatory modalities during pre-training, and a phylogenetic species embedding providing taxonomic awareness across diverse fungal clades, and expression data introduced exclusively during fine-tuning (Supplemental Figure2). Although integrating five heterogeneous modalities, the final model comprises 775,858 parameters, a scale comparable to existing single-modality benchmarks such as JEDI (∼500K parameters) and CircPCBL (∼150K parameters). Two inference modes were used throughout the evaluation. Mode A used only the genome and GTF annotation without expression data and represented the primary deployment use case for understudied fungi lacking transcriptome data. Mode B additionally incorporated host gene-expression features for within-group prediction when RNA-seq data were available. The evaluation of mycoCirc was across three complementary dimensions. The first dimension assessed gene-level circRNA prediction performance on held-out species within each taxonomic group and on completely unseen external species. The second dimension examined the contribution of the pre-training protocol and of individual modalities and training signals through a series of ablation experiments. The third dimension investigated the biological behaviour of the model across gene architectures, the junction-level prediction capability unique to mycoCirc, and mechanistic interpretability through Integrated Gradients and motif analysis.

### Held-Out Baseline Evaluation and Cross-Species Zero-Shot Generalisation

Under Mode A evaluation, mycoCirc consistently achieved AUROCs between 0.690 and 0.699 across all three taxonomic groups on the held-out test set, substantially outperforming JEDI at 0.506 to 0.566 and CircPCBL at 0.492 to 0.533 (Figure 1A). However, Mode B exhibited highly variable results, where the Filamentous group performance improved to an AUROC of 0.723, while the Ascomycete yeast and Basidiomycete yeast groups degraded to 0.453 and 0.570, respectively, indicating that expression data do not reliably transfer cross-species.

**Figure 1.**
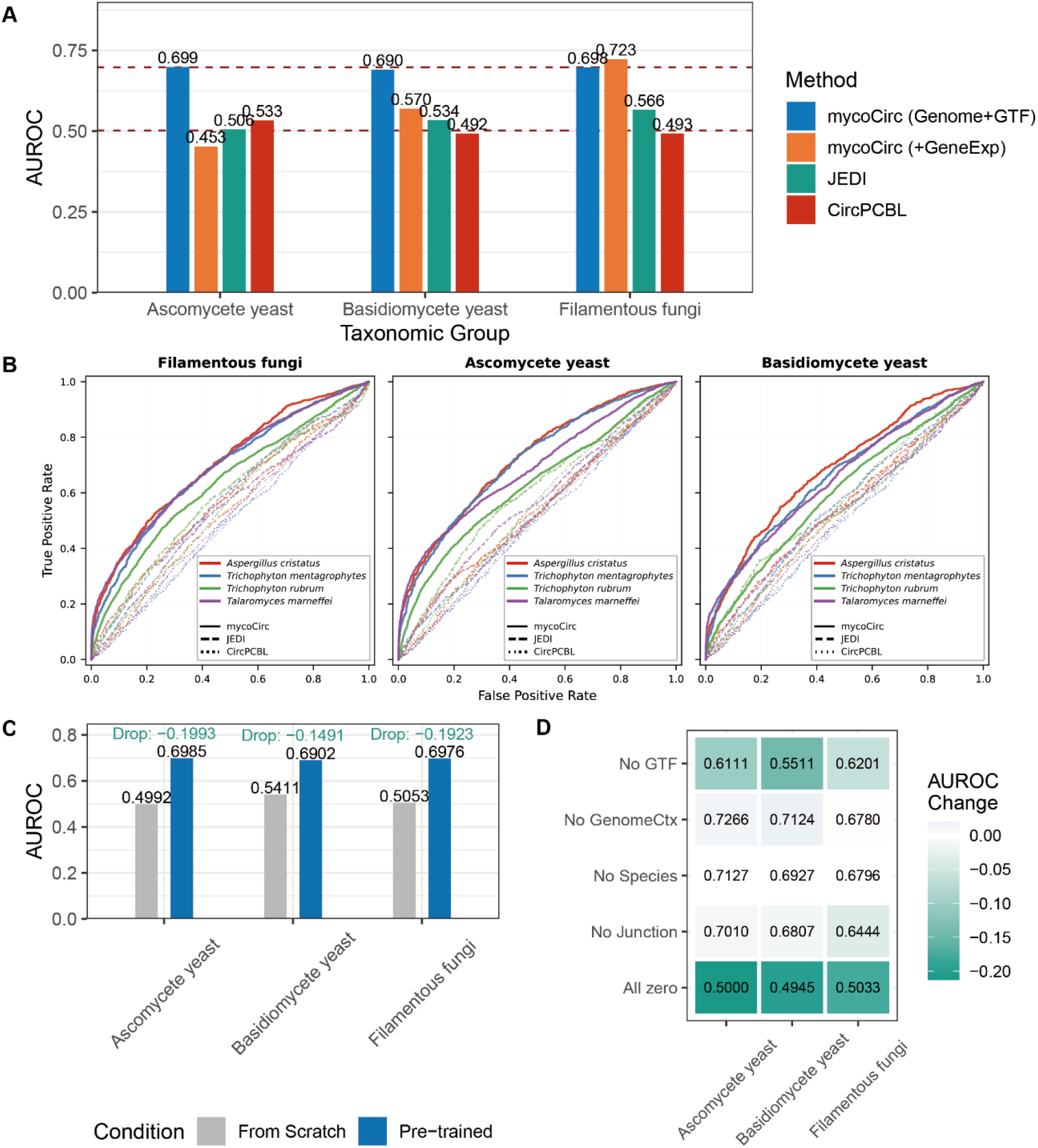
Cross-species circRNA prediction performance and model interpretation. **A. Held-out test performance**. Bar plot with groups (Candida, Cryptococcus, Filamentous) on the x-axis and AUROC on the y-axis (0-1). Each group has four bars: mycoCirc Mode A (Genome+GTF, blue), mycoCirc Mode B (Genome+GTF+GeneExp, orange), JEDI (teal), and CircPCBL (red). Dashed horizontal lines indicate AUROC = 0.5 (random classifier) and 0.7. **B. Cross-species circRNA prediction ROC curves**. ROC curves comparing mycoCirc (solid lines), JEDI (dashed lines), and CircPCBL (dotted lines) on four independent fungal species not represented in the training set. Each panel evaluates one fine-tuned models. Curves are coloured by species: Aspergillus cristatus (red), Trichophyton mentagrophytes (blue), Trichophyton rubrum (green), and Talaromyces marneffei (purple). All methods were evaluated under Mode A (genome and annotation only) using balanced test sets with matched positive and negative samples. The grey dashed diagonal represents the performance of a random classifier. **C. Pre-training ablation**. Bar plot with taxonomic groups on the x-axis and AUROC on the y-axis. Each group has two bars: from-scratch (gray) and pre-trained (blue). The average AUROC drop is annotated. **D.Component ablation**. Heatmap with removed modalities (GTF, Junction, Genome context, Species, Expression) on the x-axis and taxonomic groups on the y-axis. Colour intensity represents AUROC change relative to full Mode A (red = decrease, blue = increase), with values annotated in each cell. AUROC, area under the receiver operating characteristic curve; GTF, gene annotation file.

To further challenge the zero-shot transferability of the framework on entirely independent external datasets, we comprehensively retrieved available high-quality public total RNA-seq datasets for non-model fungi, yielding four completely unseen species encompassing *Aspergillus cristatus, Talaromyces marneffei, Trichophyton mentagrophytes, and Trichophyton rubrum* (Figure 1B and Supplemental Table 1). Utilizing Mode A, mycoCirc achieved AUROCs ranging from 0.598 to 0.722 across the twelve evaluation combinations, where the Ascomycete yeast model exhibited peak performance on *A. cristatus* at 0.722 and *T. mentagrophytes* at 0.714, while the Filamentous fungi model maximised predictions for *T. marneffei* at 0.696. This divergence reveals an uncoupling between macroscopic morphology and the genomic architecture governing circRNA biogenesis, suggesting that the Ascomycete yeast model successfully captured basal, highly conserved phylum-level features that remain predominant drivers of circularisation in certain filamentous species. Crucially, mycoCirc substantially outperformed JEDI at AUROCs of 0.525 to 0.593 and the plant-derived CircPCBL at AUROCs of 0.493 to 0.550, while the cross-phylum collapse of the basidiomycete checkpoint consistently yielded the lowest AUROCs of 0.598 to 0.686, further validating the profound impact of evolutionary distance on model transferability. For Mode B validation, where the independent *T. marneffei* cohort served as the sole unbiased external benchmark, mycoCirc yielded AUROCs of 0.703, 0.565, and 0.702 across the Ascomycete yeast, Basidiomycete yeast, and Filamentous fungi fine-tuned models, respectively. This incremental shift suggests that incorporating quantitative expression features may offer only modest gains over genomic and annotation data alone when generalising to external lineages, possibly reflecting technical variations across different sequencing pipelines.

### Systematic Ablation of Model Components and Architectural Adaptability

To quantify the driving forces behind mycoCirc, we systematically evaluated its training protocols, architectural components, and expression modalities within a unified ablation framework. First, fine-tuning the model from random initialization caused predictive performance across all three operational groups to collapse to near-random levels (AUROC 0.500–0.540, average drop: 0.180; Figure 1C), confirming that the two-stage pre-training protocol is indispensable for learning transferable cross-species representations. Second, systematic input ablation during inference revealed that the GTFEncoder serves as the primary non-expression driving force (Figure 1D), with its removal dropping the average AUROC by 0.101, whereas removing the JunctionEncoder yielded a modest drop of 0.020. This variation in scale is expected because gene-level biogenesis signals reside predominantly within macro gene-structure features captured by the GTFEncoder, whereas the JunctionEncoder specifically functions to resolve exact back-splicing junctions. Crucially, zeroing out genomic context or species embeddings had near-zero effect, indicating that local sequence composition adds minimal predictive power beyond structural annotations. Finally, while zeroing expression data triggered the sharpest performance degradation within-group (dropping by 0.157–0.204), this intra-group dominance failed to translate into cross-species predictive utility. Within-group cross-validation demonstrated that full expression data achieved an AUROC of 0.994, yet shuffling either CircExp or GeneExp individually caused negligible degradation (∼0.010), though zeroing both concurrently triggered a sharp drop (Figure 2A). This contrast confirms that transcriptomic data provides an exceptionally robust yet highly redundant intra-group signal where either abundance channel suffices. Consequently, the cross-species fragility of transcriptomic abundance highlights its high susceptibility to evolutionary drift and laboratory batch effects, establishing those structural genomic annotations offer far more stable, evolutionary invariant features for pan-fungal prediction than mutable RNA-seq quantification.

**Figure 2.**
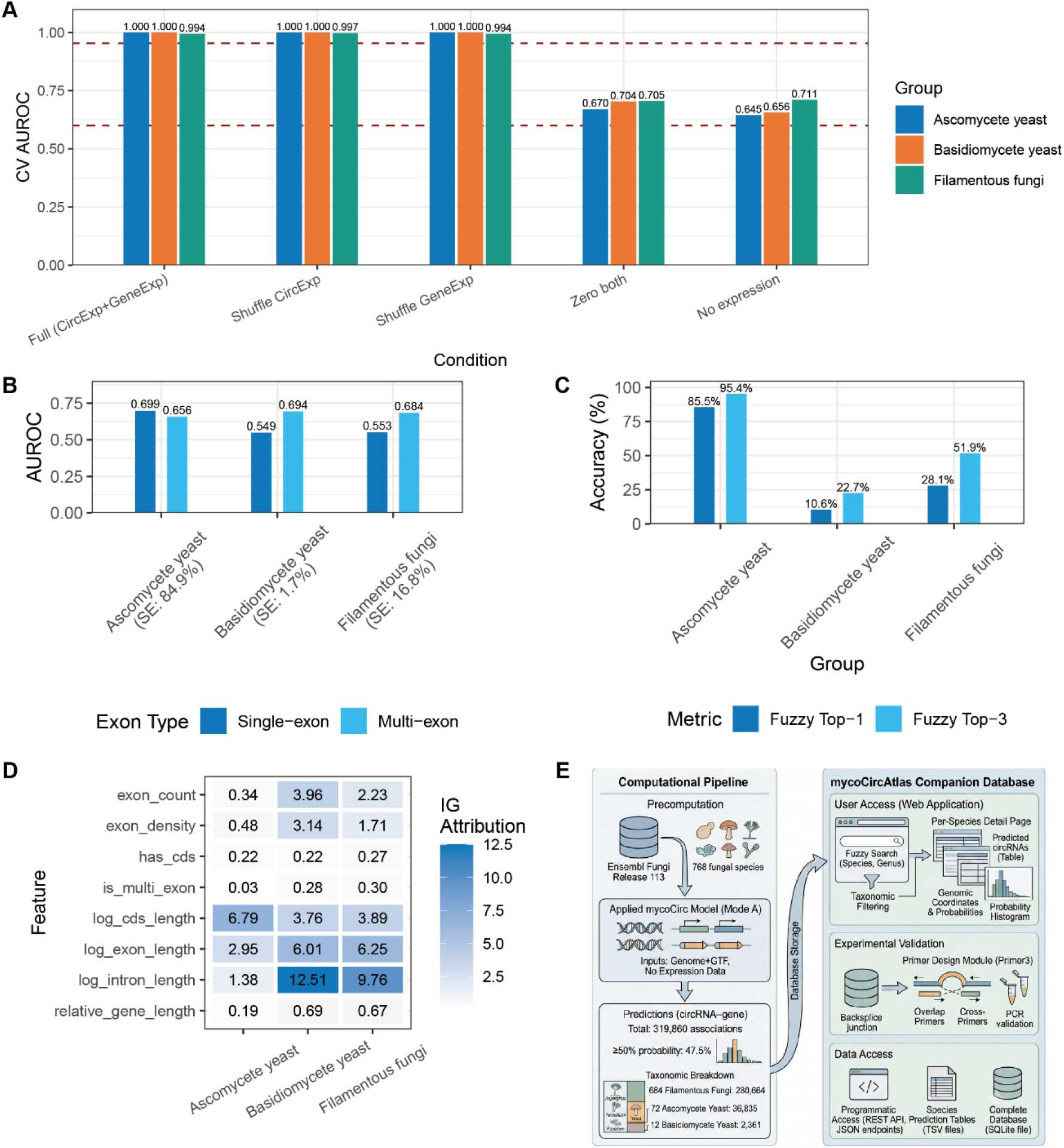
Additional analyses supporting model robustness and interpretability. **A. Expression ablation within-group cross-validation**. Bar plot with five perturbation conditions on the x-axis (Full, Shuffle CircExp, Shuffle GeneExp, Zero both, No expression) and CV AUROC on the y-axis. Three bars per condition represent Candida (blue), Cryptococcus (orange), and Filamentous (teal). **B. Exon type breakdown**. Grouped bar plot with taxonomic groups on the x-axis, AUROC on the y-axis. Each group has two bars: single-exon (blue) and multi-exon (cyan). The percentage of single-exon genes among circRNA-positive genes (SE%) is embedded in each group label on the x-axis. AUROC, area under the receiver operating characteristic curve; GTF, gene annotation file; SE, single-exon. **C. Backsplice junction prediction accuracy**. Grouped bar plot with taxonomic groups on the x-axis and accuracy (%) on the y-axis. Each group has bars for Top-1 and Top-3 fuzzy matching accuracy (allowing ±5 bp tolerance at the nucleotide level). **D. GTF feature importance**. Heatmap with GTF features (CDS length, Exon length, Intron length, Exon density, Exon count) on the y-axis and taxonomic groups on the x-axis. Colour intensity represents Integrated Gradients attribution score (higher = more important), with values annotated in each cell. **E. Overview of the mycoCircAtlas companion database and computational pipeline**. The schematic map illustrates the database framework designed to provide comprehensive precomputed gene–circRNA predictions across the fungal kingdom. AUROC, area under the receiver operating characteristic curve; CV, cross-validation; CircExp, circRNA expression; GeneExp, gene expression; GTF, gene annotation file; IG, Integrated Gradients.

Given that structural genomic annotations serve as the primary driving force for mycoCirc, we next investigated whether this structural reliance led to performance variations across distinct gene architectures, specifically comparing single-exon and multi-exon profiles (Figure 2B). In the Ascomycete yeast group, where single-exon architectures predominate (constituting 88.5% of all genes and 84.9% of circRNA-producing genes), the model achieved AUROCs of 0.699 on single-exon and 0.656 on multi-exon genes. Conversely, in the Basidiomycete yeast and Filamentous fungi groups, where single-exon genes constitute the structural minority (representing 2.40% and 21.1% of all genes, respectively), predictive performance on this minority class aligned at AUROCs of 0.549 and 0.553, while robust multi-exon prediction was maintained at 0.694 and 0.684. While the lower multi-exon performance in Ascomycete yeasts warrants a cautious interpretation due to the limited sample size of this architectural subset (35 multi-exon genes with 23 positive cases), the overall divergence across lineages remains fundamentally consistent with the GTFEncoder’s reliance on macro-structural features. Because single-exon genes lack complex intronic boundaries, they inherently present fewer discriminative spatial cues, thereby bottlenecking the framework’s structural pattern recognition in lineages where such architectures are atypical.

### Granular Back-Splice Junction Resolution and Positional Ranking Performance

Building upon macro-level gene classification, we evaluated whether mycoCirc could resolve micro-level splice sites. This capability leverages the JunctionEncoder’s cross-attention mechanism to rank all potential back-splice exon pairs within a gene. We assessed performance via Top-1 accuracy (the true junction ranks first among all candidates) and Top-3 accuracy (the true junction ranks within the top three candidates) using a fuzzy matching strategy with a ±5 bp tolerance (Figure 2C).The Ascomycete yeast group achieved 85.5% Top-1 and 95.4% Top-3 accuracy, benefiting from a compact, single-exon-dominated architecture with few candidate boundaries. Conversely, the Basidiomycete yeast group lagged at 10.6% Top-1 and 22.7% Top-3 accuracy, while the Filamentous fungi group sustained 28.2% Top-1 and 51.9% Top-3 accuracy, demonstrating moderate success in complex multi-exon contexts. This reduced precision in multi-exon-dominant taxa likely stems from sequencing-depth limits on lowly expressed junctions, alignment ambiguity at dense exon boundaries, and the mathematical difficulty of ranking under severe class imbalance.

### Mechanistic Feature Attribution and Interpretability

To elucidate the underlying molecular driving forces shaping these taxon-specific performance profiles, we applied Integrated Gradients to decipher the precise genomic features prioritized by mycoCirc. Moving beyond the macro-level dependencies identified in our input ablation, this attribution analysis revealed that within the dominant GTFEncoder modality, feature importance rankings diverged in a manner that precisely mirrored each group’s evolutionary genome configuration (Figure 2D). In the single-exon-dominated Ascomycete yeast group, coding sequence (CDS) length emerged as the premium feature, serving as a critical proxy for overall transcript architecture. Conversely, intron length dominated feature weighting in the Basidiomycete yeast group, reflecting the heightened reliance of intron-rich yeast genomes on splicing machinery to facilitate back-splicing. Similarly, in the Filamentous fungi group, intron length occupied the highest rank, closely followed by exon count and length. Despite these lineage-specific weightings, the global mechanistic patterns remained highly coherent: genes characterized by longer introns and greater exon counts offer expanded spatial opportunities for back-splicing events.

Given that flanking sequence composition modulates circularization efficiency, we extended our interpretability analysis to localized nucleotide preferences within the JunctionEncoder. *K*-mer attention mapping revealed a significant enrichment of A/T-rich motifs—specifically CAA, TTT, and AAG—in immediate proximity to back-splice junctions (Supplemental Figure 3). This localization aligns with the established A/T sequence bias in fungal introns that correlates with enhanced back-splicing efficiency. Collectively, these interpretable feature weights confirm that mycoCirc leverages the deep evolutionary blueprints governing fungal transcript processing rather than memorizing superficial training correlations.

### The mycoCircAtlas Companion Database for Fungal Functional Genomics

To maximise utility for the fungal research community, mycoCircAtlas was constructed as a companion database providing precomputed predictions across the fungal kingdom (Figure 2E). The appropriate fine-tuned mycoCirc model was applied under Mode A, utilising only genomes and structural annotations, to all 768 fungal species in Ensembl Fungi Release 113. This screening yielded 319,860 high-confidence circRNA associations, spanning 72 ascomycete yeast species (36,835 predictions), 12 basidiomycete yeast species (2,361 predictions), and 684 filamentous fungi species (280,664 predictions). The most densely represented genera included *Aspergillus, Penicillium, Fusarium*, and *Colletotrichum*, reflecting their prevalence among sequenced fungal genomes.

The database is deployed as a publicly accessible web application supporting fuzzy queries by species, genus, or common name alongside taxonomic filtering. Each species page provides a structured table detailing predicted genomic coordinates, probabilities, and exon counts. Furthermore, an integrated primer-design module automatically generates polymerase chain reaction validation primers targeting specific back-splice junctions using Primer3, outputting both overlap primers spanning the junction and cross-primers flanking the junction to facilitate downstream experimental validation.

## Discussion

The mycoCirc framework addresses a critical computational bottleneck imposed by the high cost and technical challenges of fungal transcriptomic sequencing. By evaluating this framework across phylogenetically diverse lineages, this study demonstrated that multi-modal structural encoding can overcome the limitations of sequence-only methods. The core computational driver of this integrated architecture is the GTFEncoder, which supplies critical gene-architecture context completely inaccessible to sequence-only approaches. This structural insight explains why historical sequence-only models consistently underperformed when applied to complex fungal datasets. Concurrently, the JunctionEncoder plays a complementary role within the network. Although its contribution to gene-level metrics remains modest because primary circularisation signals reside predominantly within macro-topological gene structures, its main utility is found in the junction-prediction task by enabling the precise ranking and identification of back-spliced exon pairs.

The combination of pan-fungal pretraining and lineage-specific fine-tuning enables robust zero-shot cross-species generalisation. This two-stage pretraining protocol captures conserved features of circRNA biogenesis that transcend specific generic boundaries, thereby establishing a practical predictive alternative for species entirely lacking transcriptomic resources. However, the analysis of exon types exposes a distinct biological boundary governing model performance. Within single-exon-dominated lineages, single-exon prediction performance remains stable. Conversely, within multi-exon-dominant taxa, single-exon prediction performance lags because those specific loci lack the rich structural coordinates upon which the GTFEncoder relies. This finding reflects a genuine biological reality wherein single-exon circRNA formation in fungi operates through molecular mechanisms fundamentally distinct from multi-exon back-splicing.

Several inherent challenges within fungal circRNA biology merit closer methodological consideration. Back-splice junction prediction within highly skewed label spaces remains a pervasive computational bottleneck. Although the binary cross-entropy loss functions employed in this study delivered strong baseline performance in compact genomes, a ranking-based loss function could offer a mathematically superior framework to mitigate severe class imbalances in multi-exon species. Furthermore, the attenuated performance observed for single-exon genes within multi-exon-dominant taxa stems from a fundamental biological constraint dictated by the intrinsic scarcity of distinctive structural signatures. While addressing this data sparsity remains an open challenge, tailored data augmentation strategies may offer a viable future remedy.

The performance drop observed under expression-based validation across yeast lineages highlights a critical evolutionary reality. Quantitative transcriptomic profiles lack transferable predictive signatures across deeply diverged lineages due to profound expression-level drift and laboratory batch effects, which reinforces the necessity of relying on invariant genomic blueprints. Additionally, the standard fusion architecture operates without per-gene dynamic weighting, suggesting that a transition towards advanced attention-based multimodal fusion mechanisms could enhance adaptive feature integration. During upstream data curation, the multi-software consensus pipeline was intentionally configured with strict stringency to maximise specificity and eliminate false positives. Sacrificing a portion of true positives through false-negative exclusion remained an inevitable trade-off when protecting training data integrity from the pervasive background noise inherent in non-model fungal transcriptomes.

Future efforts should leverage these conserved principles of fungal transcript processing to bridge the gap between computational prediction and functional characterisation. The companion mycoCircAtlas database and its integrated validation tool extend the utility of this framework by enabling researchers to query and validate circRNA candidates without local computational infrastructure. These consolidated resources are expected to transform resource-limited research environments, thereby accelerating functional genomics and non-coding RNA discovery across both medical and industrial mycology.

## Acknowledgements

The authors have nothing to declare.

## Code Availability

mycoCirc is freely available at https://github.com/yukkikou/mycoCirc.

## Tables

**Table 1.** Cross-species evaluation of mycoCirc, JEDI, and CircPCBL on four independent fungal species. Area under the receiver operating characteristic curve (AUROC) for mycoCirc under Mode A (Genome + GTF) and Mode B (+ RNA-seq gene expression), alongside JEDI and the pretrained CircPCBL Plant model.

| Species | Fine-tuned model | **mycoCirc Mode A** | **mycoCirc Mode B** | JEDI | CircPCBL |
| --- | --- | --- | --- | --- | --- |
| *Aspergillus cristatus* | Ascomycete yeast | **0.722** |  | 0.533 |  |
|  | Basidiomycete yeast | **0.686** |  | 0.549 | 0.524 |
|  | Filamentous fungi | **0.705** |  | 0.571 |  |
| *Trichophyton mentagrophytes* | Ascomycete yeast | **0.714** |  | 0.526 |  |
|  | Basidiomycete yeast | **0.66** |  | 0.552 | 0.493 |
|  | Filamentous fungi | **0.685** |  | 0.574 |  |
| *Trichophyton rubrum* | Ascomycete yeast | **0.627** |  | 0.593 |  |
|  | Basidiomycete yeast | **0.598** |  | 0.555 | 0.55 |
|  | Filamentous fungi | **0.634** |  | 0.579 |  |
| *Talaromyces marneffeii* | Ascomycete yeast | **0.687** | **0.703** | 0.562 |  |
|  | Basidiomycete yeast | **0.653** | **0.565** | 0.534 | 0.508 |
|  | Filamentous fungi | **0.695** | **0.702** | 0.536 |  |

**Supplemental Table 1.** Overview of Dataset Allocation and Analytical Workflows.

| Species Group | Species/<br>Strain<br>Count | Data<br>Origin | Used in<br>Upstream<br>Labeling<br>(circRNA<br>Calling)? | Used in<br>Mode A<br>Training/T<br>esting? | Used in<br>Mode B<br>Validation? |
| --- | --- | --- | --- | --- | --- |
| Ascomycete yeasts | 6 species/7 strains (In-house) | Newly sequenced | Yes | Yes (Train/Internal Test) | No (Internal only) |
| Basidiomycete yeasts | 5 species/7 strains (In-house) | Newly sequenced | Yes | Yes (Train/Internal Test) | No (Internal only) |
| Filamentous fungi | 8 species/8 strains (In-house) | Newly sequenced | Yes | Yes (Train/Internal Test) | No (Internal only) |
| External Unseen Species | 3 species (Public) | Public Database | No (External publications) | No | No (Biased for Mode B) |
|  | 1 species (Public) | Public Database | No (External publications) | No | Yes (Unbiased External Test) |

## Figures

**Supplemental Figure 1.**
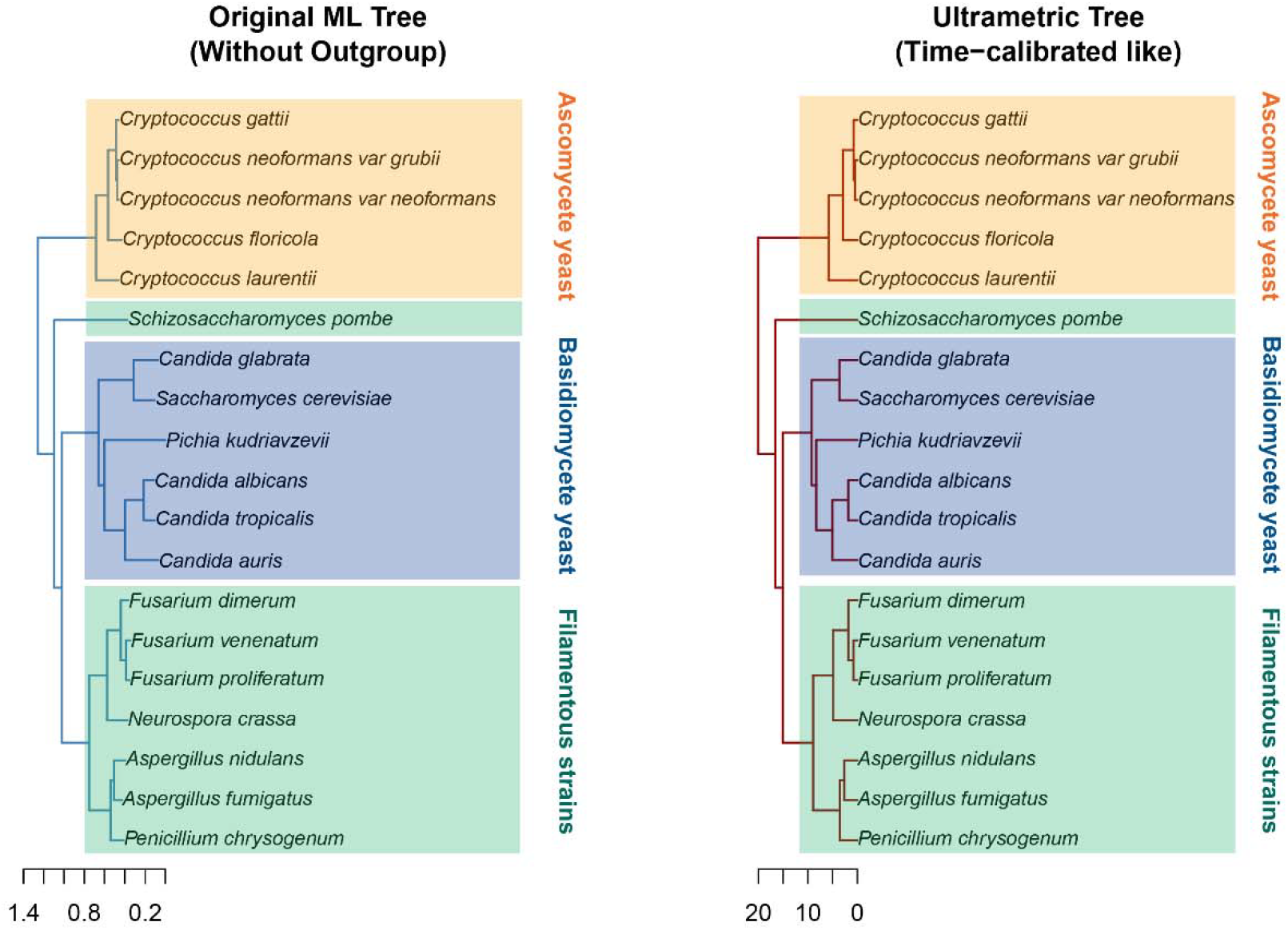
Phylogenetic landscape and evolutionary relationships across the 22 fungal strains from 19 fungi used in this study. The topology outlines the evolutionary divergence of the investigated fungal taxa, categorized into three operational functional groups. (Left) The Maximum Likelihood phylogenetic tree reveals branch lengths proportional to the estimated number of nucleotide substitutions per site. (Right) The corresponding ultrametric (time-calibrated-like) tree projects the relative divergence times, constraining all terminal nodes to an equidistant horizon from the root. Scale bars at the bottom quantify evolutionary distances adhering to the respective phylogenetic reconstruction algorithms.

**Supplemental Figure 2.**
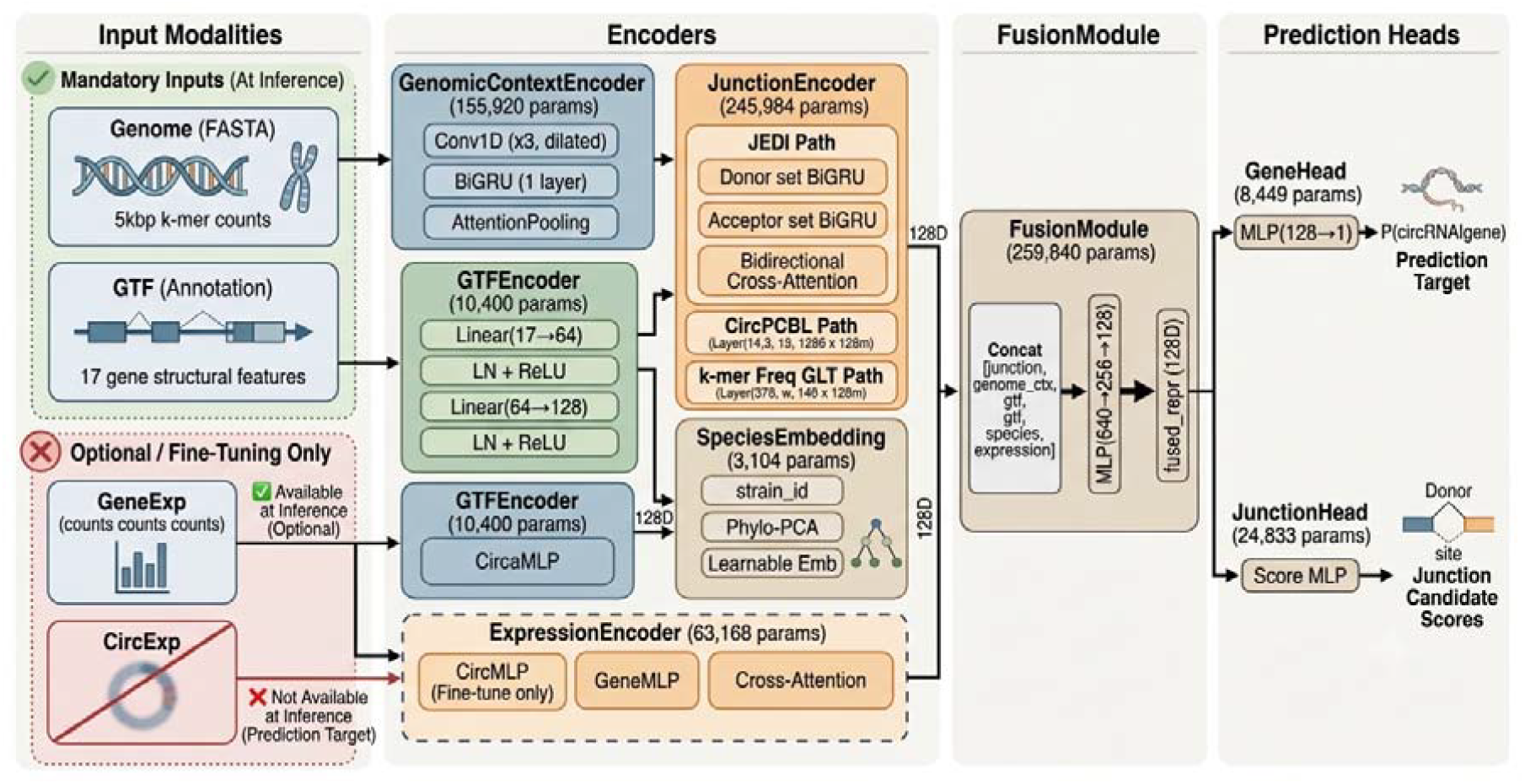
mycoCirc model architecture schematic. The model consists of specialized encoders for multi-modal inputs, a robust fusion module, and specialized prediction heads. Critically, only the Genome (FASTA) and GTF (Annotation) inputs are mandatory during the inference stage, with expression and metadata available for improved accuracy only when provided.

**Supplemental Figure 3.**
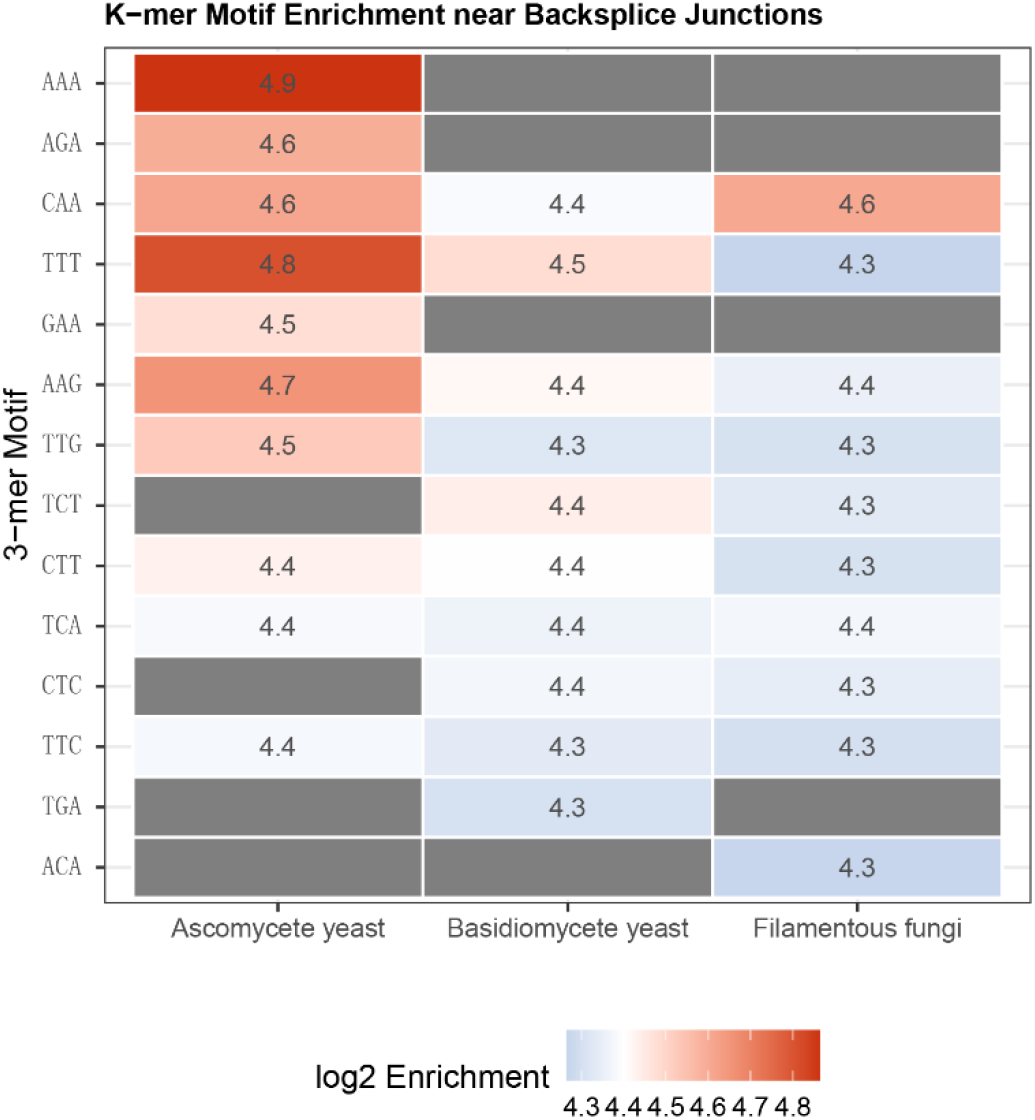
K-mer motif enrichment near backsplice junctions. Heatmap with 3-mer motifs on the y-axis and taxonomic groups (Candida, Cryptococcus, Filamentous) on the x-axis. Colour gradient from blue (low) through white to red (high) represents log2 enrichment of each motif in high-attention positions versus background sequences. Values are annotated within each cell.

